# Intersecting vulnerabilities: Climatic and demographic contributions to future population exposure to *Aedes*-borne viruses in the United States

**DOI:** 10.1101/732644

**Authors:** Guillaume Rohat, Andrew Monaghan, Mary H. Hayden, Sadie J. Ryan, Olga Wilhelmi

**Affiliations:** Institute for Environmental Sciences, University of Geneva, Switzerland; Faculty of Geo-Information Science and Earth Observation, University of Twente, Enschede, The Netherlands; National Center for Atmospheric Research (NCAR), Boulder, CO, United States; Research Computing, University of Colorado Boulder, Boulder, CO, United States; Trauma, Health and Hazards Center, University of Colorado, Colorado Springs, CO, United States; Department of Geography, University of Florida, Gainesville, FL, United States; Emerging Pathogens Institute, University of Florida, Gainesville, FL, United States; School of Life Sciences, University of KwaZulu-Natal, Durban, South Africa

## Abstract

Understanding how climate change and demographic factors may shape future population exposure to viruses such as Zika, dengue, or chikungunya, transmitted by *Aedes* mosquitoes is essential to improving public health preparedness. In this study, we combine projections of cumulative monthly *Aedes*-borne virus transmission risk with spatially explicit population projections for vulnerable demographic groups (age and economic factors) to explore future county-level population exposure across the conterminous United States. We employ a scenario matrix – combinations of climate and socioeconomic scenarios (Representative Concentration Pathways and Shared Socioeconomic Pathways) – to assess the full range of uncertainty in emissions, socioeconomic development, and demographic change. Human exposure is projected to increase under most scenarios, up to +177% at the national scale in 2080 relative to 2010. Projected exposure changes are predominantly driven by population changes in vulnerable demographic groups, although climate change is also important, particularly in the western region where future exposure may decrease by >30% under the strongest climate change mitigation scenario. The results emphasize the crucial role that socioeconomic and demographic change play in shaping future population vulnerability and exposure to *Aedes*-borne virus transmission risk in the United States, and underscore the importance of including socioeconomic scenarios in projections of climate-related vector-borne disease impacts.

## 1. Introduction

*Aedes* mosquitoes can transmit dengue, chikungunya and Zika viruses (Leta et al., 2018). The geographic range of *Aedes* mosquitoes has expanded in the conterminous United States over the past 2-3 decades (Hahn et al., 2016, Kraemer et al., 2019). Sporadic, autochthonous transmission of all three viruses has occurred recently in south Florida and Texas (Brunkard et al., 2007, Ramos et al., 2008, Trout et al., 2010, Kendrick et al., 2014, Hotez et al., 2018, Rosenberg et al., 2018). It is essential to understand how climatic and demographic changes may influence the transmission of these viruses during the 21st century.

Estimating future population exposure (i.e., the number of persons exposed to a risk of vector-borne disease transmission) to *Aedes*-borne virus transmission risk under changing climatic conditions requires an understanding of (*i*) the expansion and redistribution of *Aedes* vectors due to climate change, (*ii*) the differential vulnerability of local population groups, and (*iii*) the growth and future spatial distribution of vulnerable populations. While the influence of climate change on the expansion and redistribution of *Aedes* mosquitoes and *Aedes*-borne virus transmission risk has been explored in a wide range of studies (e.g. Campbell et al., 2015, Liu-Helmersson et al., 2019, Ryan et al., 2019), the use of projected population growth rates and patterns to estimate future population vulnerability and exposure to *Aedes* mosquitoes and *Aedes*-borne virus transmission risk is less common (Monaghan et al., 2016, Kraemer et al., 2019, Messina et al., 2019). The omission of these population projections, and lack of consideration of population subgroups, is potentially problematic in that it may lead to an overestimation of the role that climate change plays in shaping future population exposure to vector-borne diseases (VBDs) and introduces systematic bias into climate-related health adaptation planning (Ebi et al., 2016, Suk, 2016), and may lead to skewed estimates of impact across socio-demographic subgroups of the population.

In the past few years, the climate change research community has been engaged in the operationalization of a new scenario framework that facilitates the integration of future demographic and socioeconomic characteristics – through scenarios – into climate impacts, adaptation, and vulnerability (IAV) studies (Moss et al., 2010). This scenario framework is made up of climate scenarios – Representative Concentration Pathways, RCPs (van Vuuren et al., 2011) – and socioeconomic scenarios – Shared Socioeconomic Pathways, SSPs (O’Neill et al., 2015) – combined together into a scenario matrix (Ebi et al., 2013). This framework (hereafter referred as SSP*RCP framework) is being increasingly used in IAV studies to explore future population exposure – under socioeconomic and climatic uncertainty – to a wide range of climate-related risks such as extreme heat (e.g., Jones et al., 2018, Rohat et al., 2019), inland and coastal flooding (e.g., Alfieri et al., 2015, Brown et al., 2018), fire risk (Knorr et al., 2016), air pollution (Chowdhury et al., 2018), and food security (e.g., Hasegawa et al., 2014). The SSP*RCP framework has been applied to some VBD-related studies (e.g., Monaghan et al. 2016, Li et al., 2019, Messina et al., 2019). However, uncertainty in future population vulnerability and exposure to VBDs could be much more readily assessed if the SSP*RCP framework approach were applied more broadly and thoroughly across many different VBDs, particularly given the wide range of future socioeconomic pathways that exist.

In this paper, we apply the SSP*RCP framework to assess future population exposure to *Aedes*-borne virus transmission risk (VTR) in the conterminous United-States (hereafter referred as CONUS), at the county-level, up to 2080, under four consistent combinations of climate and socioeconomic scenarios. We combine projections of cumulative monthly risk of *Aedes*-borne virus transmission (under two climate scenarios) with population projections for a number of vulnerable demographic groups (under three socioeconomic/demographic scenarios). Using a scenario matrix, we explore separately the relative contribution of climate change and demographic growth to future exposure, and assess the avoided exposure due to strong mitigation options or to different socioeconomic pathways.

## 2. Data and Methods

### 2.1. Scenario setting

We explored future population exposure to *Aedes*-borne VTR under several climate and socioeconomic scenarios, spanning the wide range of uncertainties in future emission levels, socioeconomic development, and demographic growth. We employed the lowest and highest emission scenarios, RCP2.6 and RCP8.5. The former assumes strong mitigation options and a rapid decline in emissions by the middle of the century, while the latter depicts continuing business-as-usual emissions throughout the century (van Vuuren et al., 2011). The projected increase in global average temperature for 2081-2100 ranges from 0.3°–1.7°C under RCP2.6 to 2.6°–4.8°C under RCP8.5, relative to 1986-2005 (IPCC, 2013).

We combined these two emission scenarios with three socioeconomic/demographic scenarios – SSP1, SSP3, and SSP5 – covering the full range of uncertainty in demographic growth in the United States (Figure S1). Along with assumptions of population growth among different demographic groups, these scenarios also depict varying levels of socioeconomic development in terms of economic growth, environmental awareness, education, spatial patterns of urban development, technological development, health equity, and economic inequalities (O’Neill et al., 2015). SSP1, named *Sustainability*, depicts medium population growth in the United States, along with economic development that places large emphasis on human well-being and achieving development goals, reducing inequality, concentrating urbanization, and increasing sustainable consumption. By contrast, SSP3, named *Regional Rivalry*, depicts overall population decline in the United States, along with increased inequality, reduced health and education investments, slowing global economic growth, and strong governmental focus on regional security with a subsequent reduction in immigration. Finally, SSP5, named *Fossil-fueled Development*, depicts high population growth in the United States driven primarily by immigration, along with a high technological development, strong investments to enhance human and social capital, and a rapid growth of the global economy through heavy use of fossil fuel resources.

Although a given RCP can be consistent with several SSPs, not all SSP*RCP combinations are consistent, and some require more mitigation efforts than others (Kriegler et al., 2014, Kriegler et al., 2012). SSP1 and SSP5 can theoretically lead to emission levels depicted under RCP2.6 (requiring massive mitigation efforts under SSP5), but this is not the case for SSP3 (Rogelj et al., 2018). Similarly, the socioeconomic development depicted under SSP1 is not consistent with the high emission levels associated with RCP8.5. Bearing in mind these implausible combinations, we employed the SSP*RCP combinations depicted in Table 1. To enable isolating the individual contribution of climate change and population growth on future human exposure, we also explored future population exposure under combinations of (*i*) current climate and future population and (*ii*) current population and future climate (see section 2.2).

**Table 1.** Combinations of climate and socioeconomic scenarios to explore future population exposure and to isolate the climate and population (pop.) effects (see section 2.2)

|  | Current | SSP1 | SSP3 | SSP5 |
| --- | --- | --- | --- | --- |
| Current | *Baseline* | *Pop. effect* | *Pop. effect* | *Pop. effect* |
| RCP2.6 | *Climate effect* | X |  | X |
| RCP8.5 | *Climate effect* |  | X | X |

### 2.2. Exposure projections, individual effects, and avoided exposure

We defined the population exposure in a given county and for a given population group as being the combination of the cumulative monthly transmission risk of *Aedes*-borne virus with the population count. Population exposure is therefore expressed in terms of person-months of exposure per year, in line with metrics used in other climate impact studies, e.g. (Jones et al., 2015, Rohat et al., 2019). The main advantage of this exposure metric lies in that it accounts for the duration (in months) of the exposure event. We assessed current (year 2010) and future (years “2050” and “2080”) population exposure to *Aedes*-borne VTR for different population groups separately (see Section 2.4), under the four SSP*RCP combinations detailed in Section 2.1. Using the scenario matrix and combinations with current climate or current population (Table 1), we isolated the population and climate effects. The population effect represents the changes in population exposure due to changes in population growth (as a function of demographic/socioeconomic conditions) only, while the climate effect represents the changes in population exposure due to climate change only (Jones et al., 2015). We also computed the interaction effect between the two, that is, the difference between the total projected change in exposure and the sum of the climate and population effects. The interaction effect is interesting in that it represents the process by which concurrent changes in population and climatic conditions affect the population exposure (Rohat et al., 2019). We explored the population, climate, and interaction effects at the county scale for the four SSP*RCP combinations separately for both increased and decreased exposure.

Finally, we estimated the relative avoided exposure due (*i*) to shifts in climatic conditions, that is, a shift from a high (RCP8.5) to a low (RCP2.6) emission scenario (using current population conditions), and (*ii*) to shifts in population growth patterns due to socioeconomic/demographic conditions, that is, a shift from a high (SSP5) to a medium (SSP1) population growth scenario or a shift from a high (SSP5) to a low (SSP3) population growth scenario (using current climatic conditions).

### *2.3. Aedes*-borne virus transmission risk (VTR)

We retrieved projections of *Aedes*-borne virus transmission risk (VTR) from Ryan et al. (2019), for the current period (year 2010) as well as for future time-periods – “2050” (2040-2069) and “2080” (2070-2099) – under both RCP2.6 and RCP8.5. Briefly, Ryan et al. (2019) employed a temperature-based empirically parametrized model of viral transmission (by the vectors *Ae. aegypti* and *Ae. albopictus*) along with temperature projections from four general circulation models (GCMs, see Table S1) to estimate future cumulative monthly transmission risk on a 1/12° spatial grid. The temperature bounds suitable for virus transmission (with a probability of temperature suitability > 97.5%) are 21.3—34.0°C for *Ae. aegypti* and 19.9—29.4°C for *Ae. albopictus* (see Ryan et al. (2019) for full details of the modelling approach). Here we used the projections of cumulative monthly transmission risk performed with the four GCMs and aggregated them to the county scale (using area-weighted mean) for each time-period and RCP. We employed the multi-model ensemble mean to explore future transmission risk and the interquartile range (IQR) to display inter-model uncertainties.

### 2.4. Selection and projection of vulnerable population groups

Because some population groups are more vulnerable to *Aedes*-borne diseases than others (Beard et al., 2016), we assessed future exposure for a number of potentially vulnerable population groups, in addition to the exposure of the whole population. Population groups with higher vulnerability to *Aedes*-borne diseases include those who are more likely to be bitten by *Aedes* mosquitoes and those who are more likely to suffer adverse health conditions if infected by *Aedes*-borne viruses. *Aedes* mosquitoes are primarily daytime biters and prefer to take blood meals after sunrise and in late afternoon, although at least one study has shown that they will bite in the evening under artificial lights (Chadee & Martinez, 2000). Groups more likely to be exposed to *Aedes* bites include children (more likely to play outside) and outdoor workers (Bennett & McMichael, 2010, Schulte et al., 2016). Those who tend to have homes that are more permeable (e.g., open windows instead of air conditioning and broken window screens) are also more likely to receive mosquito bites (Radke et al., 2012, Reiter et al., 2003). In this regard, low-income communities appear especially vulnerable, as they are less likely to possess and/or to use air conditioning (Hernández & Bird, 2010). Many of these at-risk communities are located in the United States-Mexico (US-MX) border region. For example, Brownsville, TX, a community located at the US-MX border, has seen sporadic transmission of *Aedes*-borne viruses. A dengue outbreak investigation in 2005 determined that 85% of the population had air-conditioning while 61% reported screens on windows and doors (Ramos et al., 2008). *Aedes* mosquitoes thrive in urban environments (Messina et al., 2016) and typically oviposit in artificial, water-filled containers (Hiscox et al., 2013); this is particularly true for *Ae. aegypti*, but also to a lesser extent for *Ae. albopictus* (Roche et al., 2015). Additionally, *Ae. albopictus*, like *Ae. aegypti* exhibits highly anthropophilic biting behavior (Delatte et al., 2009). Urban populations are, therefore, considered more vulnerable than rural ones, as urbanites are more likely to be in contact with *Aedes* mosquitoes, increasing the potential for virus transmission (Salje et al., 2019). Finally, the elderly are more likely to suffer adverse health effects if infected by *Aedes*-borne viruses (Brien et al., 2009; Dye, 2014; Badawi et al., 2018), hence making this group highly vulnerable.

#### 2.4.1. Total population, elderly, and children

Population projections at the county level in the United States were retrieved from (Hauer, 2019), who used the Hamilton-Perry method (Swanson et al., 2010) to project age-sex-race/ethnicity (ASRE) cohorts up to 2100 under the five SSPs. We retrieved projections for all ASRE cohorts (*i.e.*, the total population), for elderly (ASRE cohorts older than 65 years), and for children (ASRE cohorts comprised between 5-14 years).

#### 2.4.2. Urban population

We retrieved spatial population projections under the SSPs from (Gao, 2017), who downscaled to a 1/100° grid the 1/8° spatial projections of (Jones & O’Neill, 2016). This set of projections differentiates the urban and rural populations and accounts for SSP-specific assumptions of urban development. Using these projections, we computed the share of urban population (over the total population) at the county-level under each SSP and each 10-year period from 2010 to 2080. We then combined the SSP-, time-, and county-specific shares of urban population with the county-level population projections retrieved from Hauer (2019), yielding county-level projections of urban population under each SSP.

#### 2.4.3. Outdoor workers

We considered outdoor workers as those people who have occupations in which >70% of the work performed is outside, according to the Bureau of Labor Statistics (see Table S2; BLS, 2016). We retrieved county-level data on occupation of the employed population from the American Community Survey (ACS) estimates, spanning yearly from 2010 to 2017. We then computed the ratio of outdoor workers over the working age population (20-64 years) for each county, averaged across the period 2010-2017. This ratio ranges from 8.7% to 48.0%, with most counties being close to the national average ratio of ∼23% (Figure S2). Assuming constant county-specific ratios, we applied the population projections of 20-64 years ASRE cohorts – retrieved from Hauer (2019) – to project the future number of outdoor workers at the county-level under each SSP.

#### 2.4.4. Low-income population

We retrieved national-scale projections of population in poverty under each SSP from Rao et al. (2018), which were generated by combining Gini projections with GDP and population projections, assuming lognormal income distributions. We combined these projections with SSP-based national population projections (KC & Lutz, 2014) to estimate SSP-specific compound annual growth rates (AGRs) of poverty reduction for each 10-year step spanning 2010-2080. Assuming changes in poverty rates to be homogeneously spread across the country, we applied the SSP- and time-specific national AGRs of poverty reduction to the current county-level shares of low-income population (that is, below the national poverty threshold; data retrieved for different age-sex cohorts from the ACS estimates for year 2012) and employed the ASRE population projections retrieved from Hauer (2019) to estimate future low-income populations at the county-level under each SSP.

## 3. Results and discussion

### 3.1. Population projections

The total population of the conterminous United-States (CONUS) is projected to shift from approximately 301 million (M) in 2010 to 472 M in 2050 and to 627 M in 2080 under SSP5, plateau at 465 M in 2080 under SSP1, and decrease to 298 M in 2080 under SSP3 (Figure 1). The urban CONUS population shows similar trends, with a slightly higher growth rate compared to total population, due to the increased urbanization depicted under all the SSPs (Jiang & O’Neill, 2015). In contrast, the increase in outdoor population is slower than that of the total population, because of the relatively lower increase in working age population. Nevertheless, the number of outdoor workers still largely increases under SSP1 and SSP5, shifting respectively from 35 M in 2010 to 41 M and 58 M in 2080 (Figure 1).

**Figure 1.**
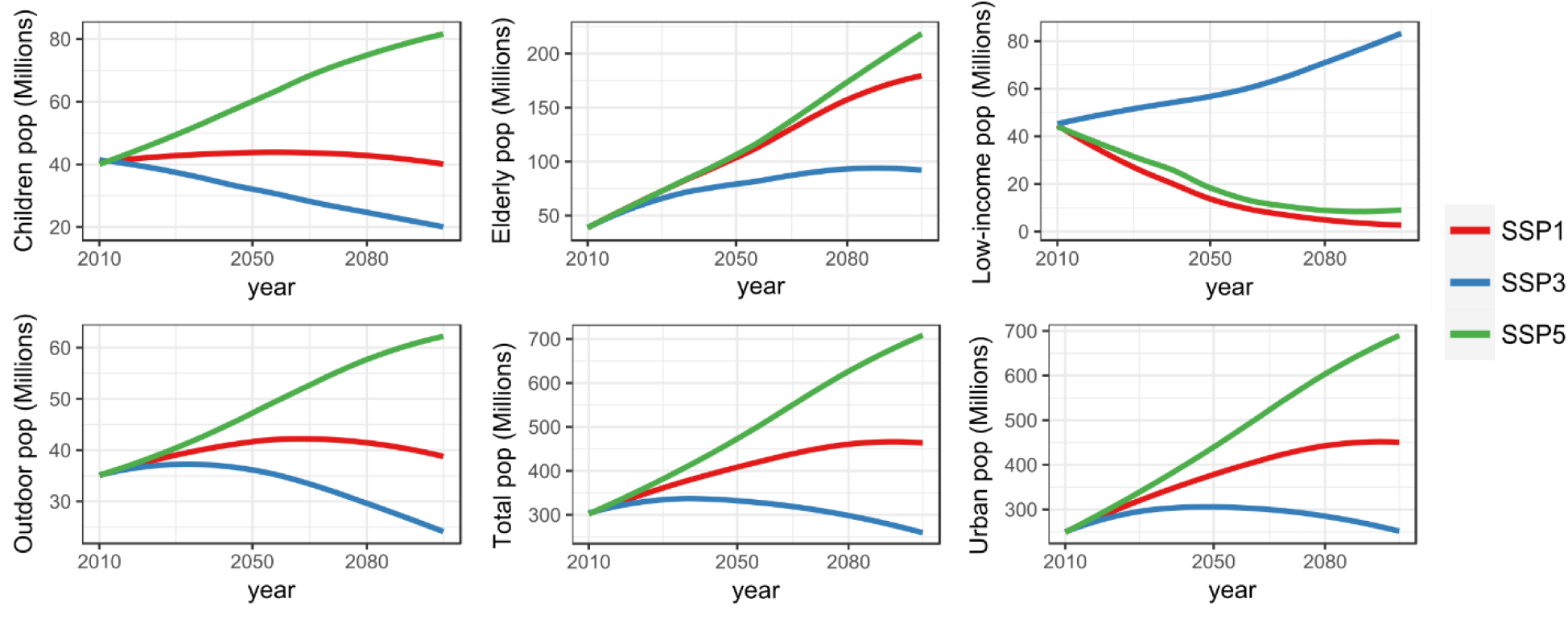
Population projections of the total population and the five vulnerable population groups, under SSP1, SSP3, and SSP5, for the conterminous United-States.

Consistent with recent trends, all SSPs (including SSP3) depict an increase in the number of elderly (older than 65 years). Noteworthy, the increase in elderly under SSP1 and SSP5 follows a similar trend, both shifting from approximately 39 M in 2010 to 105 M in 2050, and reaching 158 M (174 M) under SSP1 (SSP5) by 2080, that is, a ∼5-fold increase compared to 2010. Conversely, the ageing of the society leads to a progressive decrease in the number of children under SSP1 – and to a rapid decrease under SSP3, linked to the total population decline. The number of children increases only under SSP5, due to the high immigration-driven demographic growth.

Finally, the low-income population decreases under SSP1 and SSP5 – due to economic growth, enhancement of social capital, and strong decrease in economic inequalities –, shifting from 45 M currently to 5 M (9 M) in 2080 under SSP1 (SSP5). In contrast, SSP3 depicts an increase in the net number of low-income population – despite the decline of the total population – reaching 71 M in 2080, mainly due to the progressive decline in social welfare programs, long-term economic downturn, and increased economic inequalities.

Spatial patterns of population projections indicate great variations across regions (Figure S3) and counties (Figure 2). Despite the high demographic growth depicted under SSP5, a number of counties – predominantly located in the Midwest and South – have a declining population. SSP1 also leads to very contrasting spatial patterns, with some regions (such as Florida, California, and southern Texas) showing great population growth (> +50% in 2080 relative to 2010), while a number of counties in the Midwest and South show a large population decline of -25% to more than -50% in 2080 (relative to 2010). Noteworthy, some counties that have been rapidly growing in the past decades still show a high population growth under SSP3, despite the overall decline of the population. Altogether, the contrasting trends and spatial patterns of population projections of the vulnerable groups are likely to influence future levels and spatial patterns of population exposure.

**Figure 2.**
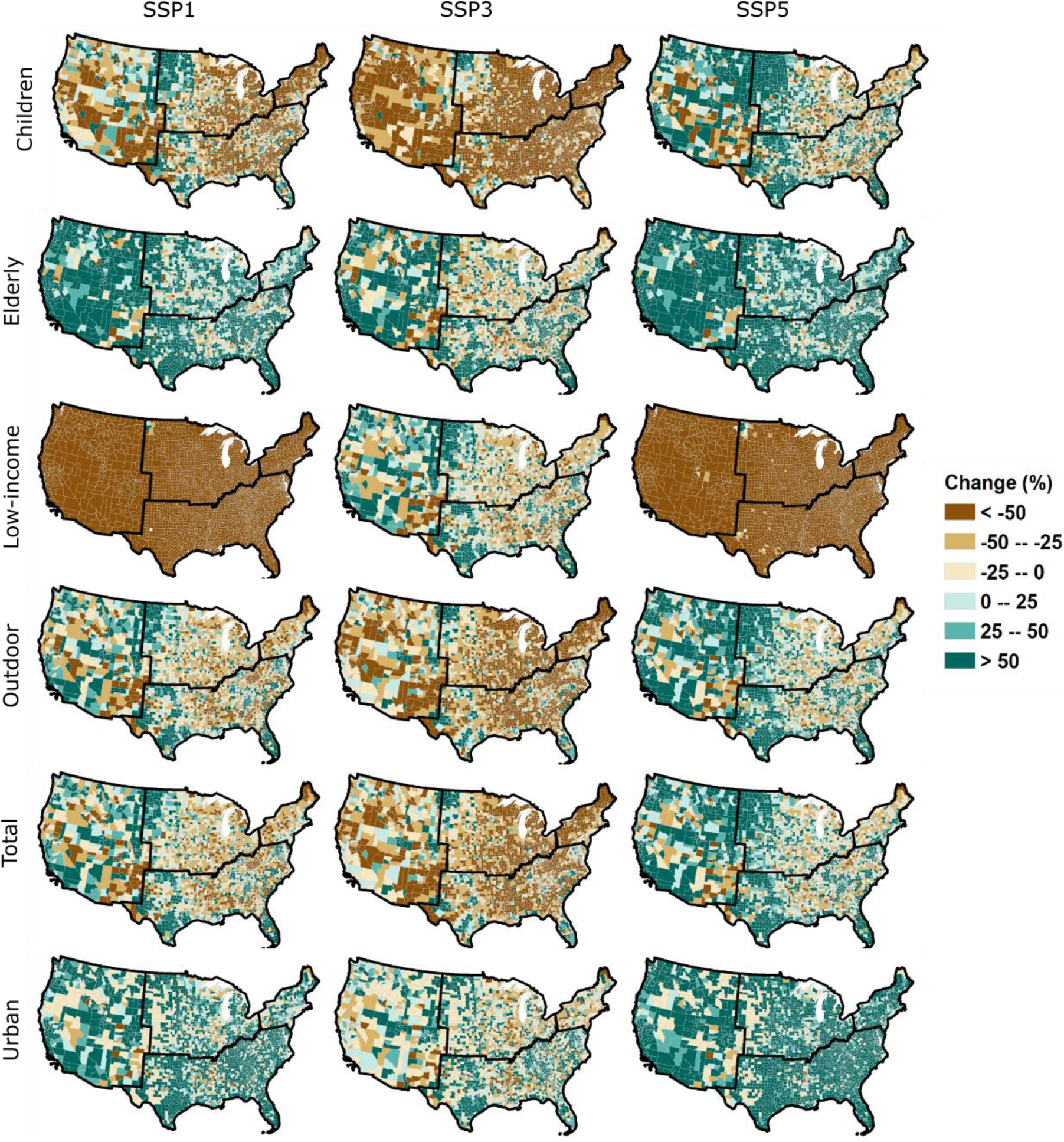
County-level spatial patterns of change in population (for year 2080 relative to year 2010), for the different population groups, under SSP1, SSP3, and SSP5.

### 3.2. Projections of cumulative monthly transmission risk of *Aedes*-borne virus

At the national level (CONUS) a significant increase in temperature suitability for VTR by the vector *Ae. aegypti* is projected under RCP8.5, with the multi-model spatial average cumulative monthly transmission risk shifting from approximately 2.8 months in 2010 to 3.5 (*IQR=0.3*) months in 2050 and 4.0 (*0.1*) months in 2080 (Figure 3a and Table S3). Under this stronger climate change scenario, some southern counties attain year-round transmission risk in 2080, while the maximum cumulative monthly transmission risk in 2010 is less than 10 months. Noteworthy, RCP8.5 leads to a much smaller increase in temperature suitability for VTR by the vector *Ae. albopictus*, with the CONUS-averaged cumulative monthly transmission risk shifting from 3.1 months in 2010 to 3.4 (*0.2*) months in 2050 and 2080. This is due to the comparatively lower maximum temperature threshold of this species (29.4°C for *Ae. albopictus* compared to 34.0°C for *Ae. aegypti*) that is increasingly exceeded under the RCP8.5 scenario, particularly in the South. In contrast, climate change as depicted under the RCP2.6 scenario has little influence on the CONUS-averaged cumulative monthly VTR by *Aedes* mosquitoes, leading only to a slight increase (0.1 month) for both vectors (Table S3).

**Figure 3.**
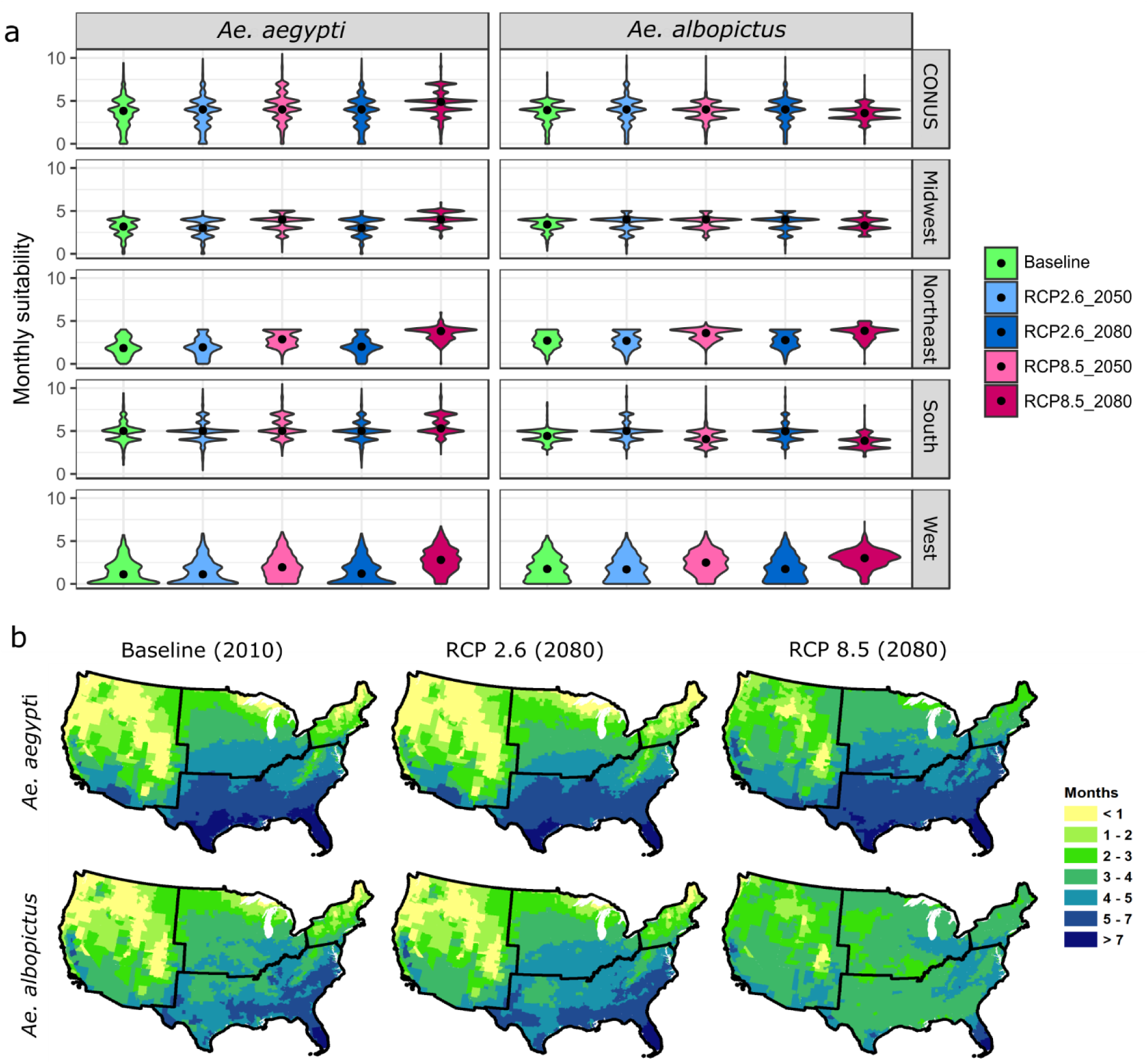
Multi-model averaged cumulative monthly VTR by *Ae. aegypti* and *Ae. albopictus*, projected under RCP2.6 and RCP8.5, represented as **(a)** the national and regional distribution of county-level results for years 2010, 2050, and 2080; and as **(b)** county-level maps for years 2010 and 2080.

The CONUS-averaged results exhibit large regional disparities (Figure 3b). The increase in cumulative monthly VTR by *Ae. aegypti* due to climate change under RCP8.5 is particularly reinforced in the West and Northeast, where it doubles in 2080, relative to 2010. In the Midwest, all counties will show suitable temperatures for virus transmission in 2080, as the minimum cumulative monthly transmission risk is 2.0 (*0.1*) months under this scenario, compared to 0 month in 2010. The number of counties in the West showing no temperature suitability year-round also largely decreases under this scenario (Figure 3b). For *Ae. albopictus*, RCP2.6 scenario leads to a significant increase cumulative monthly VTR in certain areas of the South, with values in the most at-risk counties shifting from 8.3 months in 2010 to 11.0 (*0.6*) months in 2080. Under RCP8.5, VTR decreases significantly in the South (from 4.4 to 3.8 (*0.4*) months in average in 2080 relative to 2010), but increases in the West (from 1.8 to 2.9 (*0.1*) months in average).

### 3.3. Future population exposure

Aggregated at the CONUS scale, results show an increase in total population exposure to *Ae. aegypti* VTR under all scenario combinations (Figure 4), shifting from approximately 1.14 billion (B) person-months per year in 2010 to 1.50 (*IQR=0.01*) B under SSP3*RCP8.5, 1.90 (*0.16*) B under SSP1*RCP2.6, 2.58 (*0.22*) B under SSP5*RCP2.6, and up to 3.16 (*0.03*) B under SSP5*RCP8.5 by 2080, *i.e.*, an increase in exposure ranging from 32% to 177% the current level. In comparison, the increase in total population exposure to *Ae. albopictus* VTR is lesser, shifting from approximately 1.15 B person-months in 2010 to 1.92 (*0.007*) B under SSP1*RCP2.6 and 2.61 (*0.009*) B under SSP5*RCP2.6, *i.e.*, an increase of 127% the current level. Noteworthy, total population exposure to *Ae. albopictus* VTR (*i*) remains stable at 1.15 (*0.09*) B person-months under SSP3*RCP8.5 and (*ii*) is greater under SSP5*RCP2.6 (2.61 (*0.009*) B) than under SSP5*RCP8.5 (2.42 (*0.19*) B), because of the restricting effect of a comparatively stronger climate change on temperature suitability for VTR by *Ae. albopictus* under RCP8.5.

**Figure 4.**
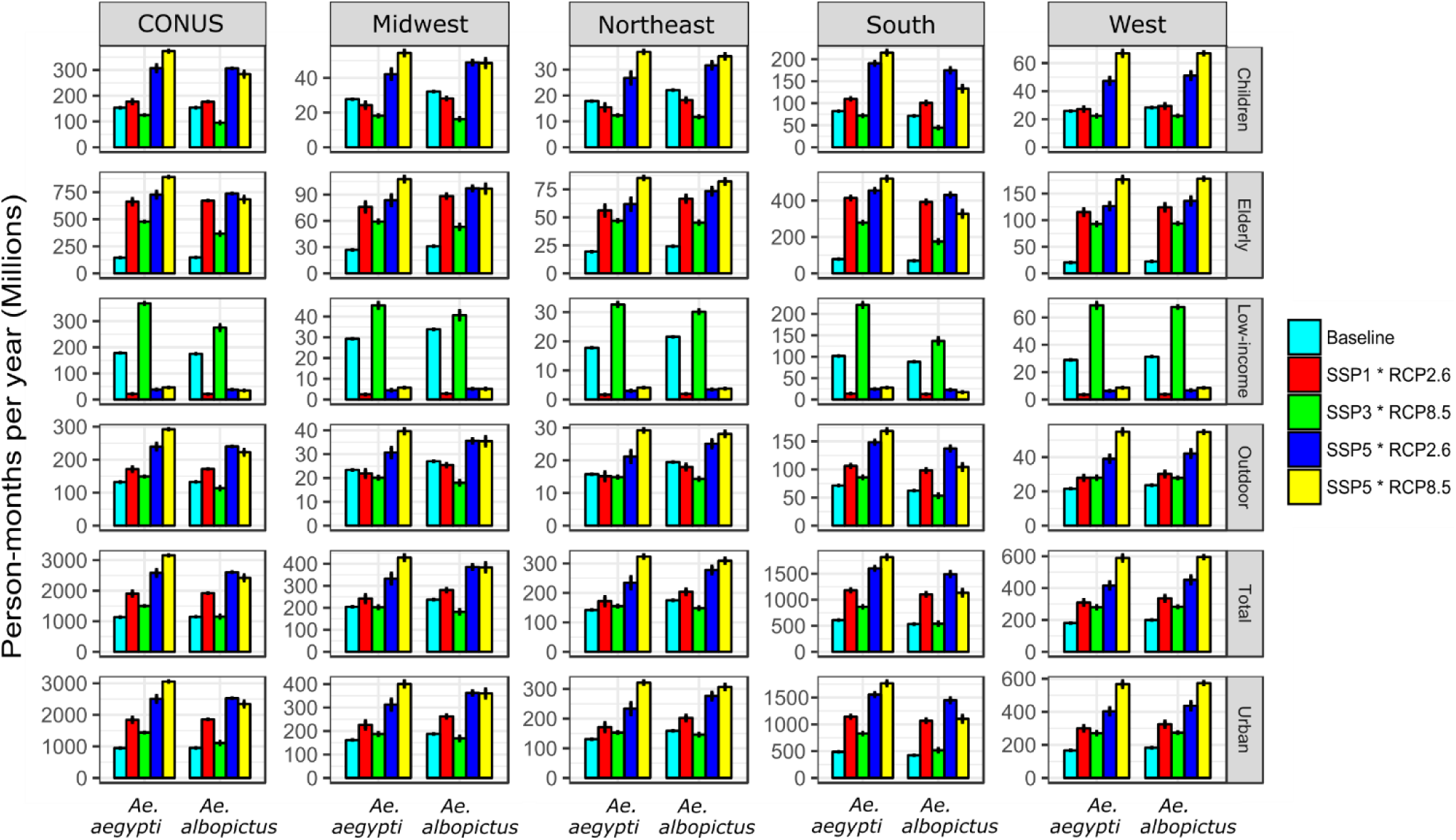
Multi-model projections of population exposure (in millions of person-months per year) to *Aedes*-borne VTR, aggregated at the continental (CONUS) and regional scale, for the current period (year 2010, Baseline) and for 2080 under four SSP*RCP combinations. Results are presented separately for the different population groups and the two *Aedes* mosquitoes. Errors bars represent the multi-model interquartile ranges (IQRs).

Not all vulnerable population groups follow similar trends in population exposure to that of the total population. Because of continuing urbanization, the increase in exposure of urban dwellers occurs slightly faster than that of the total population. Because of the ageing population depicted under all demographic/socioeconomic scenarios, the population exposure of elderly to *Aedes*-borne VTR drastically increases under all scenario combinations. Exposure of this vulnerable group to *Ae. aegypti* increases by 230% (under SSP3*RCP8.5, 478 (*5.4*) million (M) person-months) up to 514% under SSP5*RCP8.5 (890 (*11*) M) by 2080, relative to 2010 (145 M). Conversely, the exposure of children increases only slightly under SSP1*RCP2.6 and significantly decreases under SSP3*RCP8.5 – but still largely increases under SSP5*RCP2.6 and SSP5*RCP8.5 due to the high demographic growth of this population group under SSP5. Finally, the number of low-income communities exposed to transmission risk by both vectors is expected to decrease drastically under SSP1*RCP2.6, SSP5*RCP2.6, and SSP5*RCP8.5, mainly due to the decrease in the net low-income population under these two socioeconomic scenarios. In contrast, due to the increase of low-income populations depicted under SSP3, the exposure of this vulnerable group increases under SSP3*RCP8.5 and reaches 368 (*6.1*) M person-months in 2080 (for the vector *Ae. aegypti*). In comparison, this figure shrinks down to 21 (*1.7*) M person-months under SSP1*RCP2.6, highlighting the crucial role that socioeconomic pathways play in shaping future exposure.

In absolute numbers, the South is where the majority of exposure is located, accounting for 50-85% of continental exposure to *Ae. ageypti* VTR and for 46-64% (depending on time period, scenario combination, and population group) of continental exposure to *Ae. albopictus* VTR. However, the largest increase in population exposure is projected in the West, with (for instance) a total population exposure to *Ae. aegypti* shifting from approximately 181 M person-months currently to 589 (*39*) M under SSP5*RCP8.5 in 2080, which represents a 225% increase relative to 2010 (as opposed to the 177% increase at the CONUS level). The West and Northeast are the only regions where SSP5*RCP8.5 leads to a greater exposure to *Ae. albopictus* VTR than that under SSP5*RCP2.6, due to the higher temperature suitability for *Ae. albopictus* under RCP8.5 in these regions. Finally, the West and Northeast are also where the difference in exposure to *Ae. aegypti* VTR between SSP5*RCP8.5 and SSP5*RCP2.6 is the highest (regardless of the population group), suggesting a stronger influence of climate change in these regions.

### 3.4. Climate, population, and interaction effects

County-level spatial patterns of dominant effect (*i.e.*, the effect responsible for the major part of the increase or decrease in exposure) show that the population effect is the dominant contributor to both increases and decreases in total population exposure to *Ae. aegypti* VTR under SSP1*RCP2.6 and SSP5*RCP2.6 (Figure 5a). Under SSP5*RCP8.5, increases in total population exposure in counties in the West and Northeast are predominantly driven by the climate effect (*i.e.*, by climate change). Noteworthy, under SSP3*RCP8.5, the climate effect dominates the increase in total population exposure in the overwhelming majority of counties, mainly due to (*i*) decreased total population and (*ii*) stronger climate change. Results for total population exposure to *Ae. albopictus* VTR show similar trends (Figure S4), with the notable exception that the climate effect dominates the decrease in exposure in most counties of the South and Midwest under SSP3*RCP8.5 and SSP5*RCP8.5, due to the decrease in temperature suitability for *Ae. albopictus* forecasted in these regions under RCP8.5.

**Figure 5.**
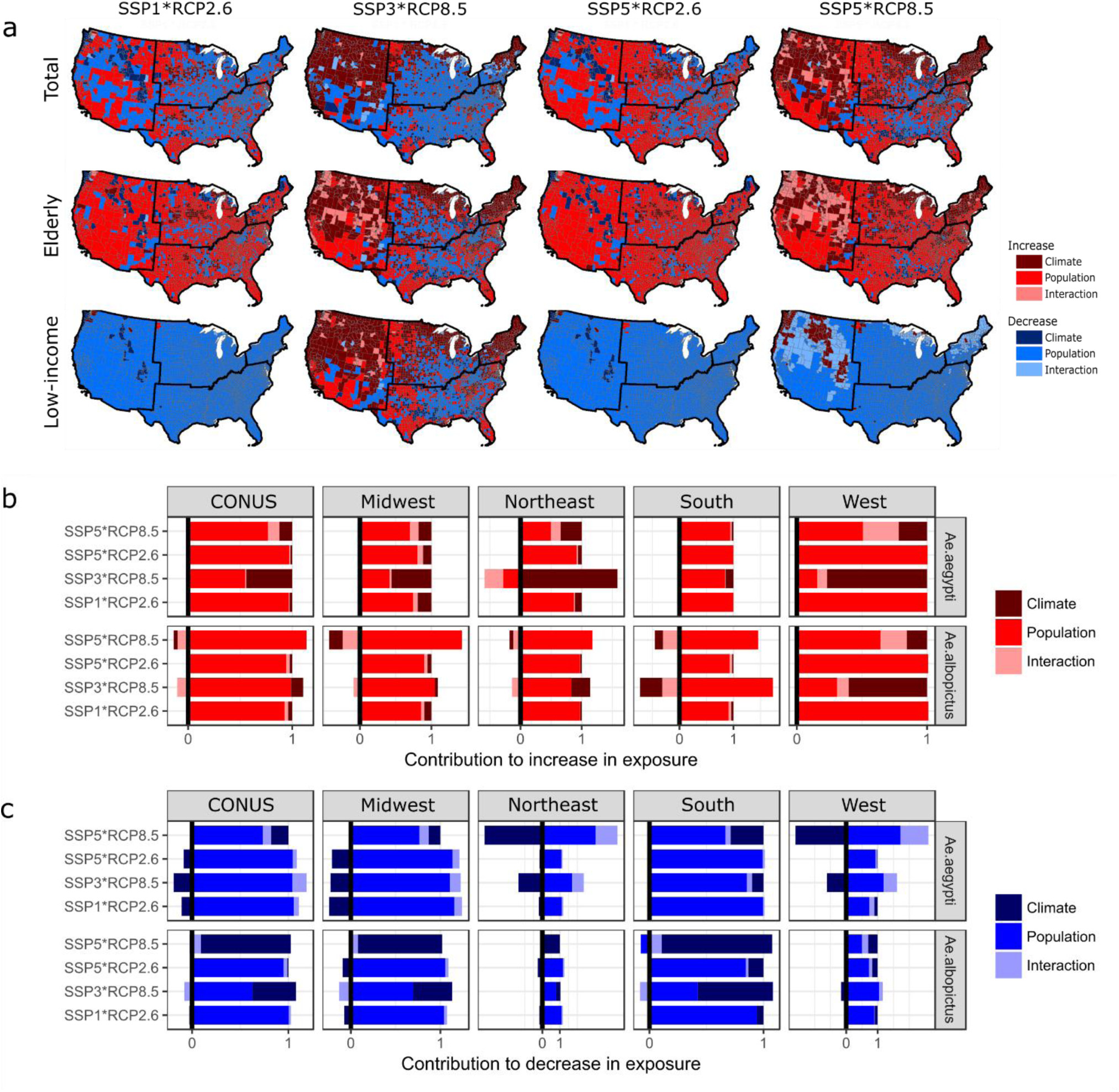
**(a)** Dominant effect (climate, population, or interaction) responsible for the highest increase (or decrease) in exposure at the county-level, for three population groups (see Figure S5 for other population groups) and for exposure to *Ae. aegypti* VTR only (see Figure S4 for exposure to *Ae. albopictus* VTR); **(b)** Contribution to increase in total population exposure of each individual effect, aggregated at the country (CONUS) and regional scale, and **(c)** same for decrease in exposure (see Figures S6-S8 for results associated with other population groups). Results are presented for year 2080 only, using the multi-model mean.

While spatial patterns of dominant effects for exposure of outdoor workers, urban population, and children are rather similar to those of the total population exposure (Figure S5), spatial patterns for the elderly and low-income communities show large differences. Results show that the population effect dominates the increase in elderly exposure to both *Ae. aegypti* (Figure 5a) and *Ae. albopictus* (Figure S4) in most counties, under all combinations (with the exception of SSP3*RCP8.5, where the climate effect dominates in many counties due to the slower growth of elderly population depicted under SSP3). Noteworthy, the interaction effect dominates the increase in elderly exposure to VTR by both *Aedes* mosquitoes in the West, highlighting the simultaneous increase in temperature suitability and growth of elderly population. Because of the strong decrease in the net number of low-income persons under SSP1 and SSP5, the population effect is the overwhelming contributor to the decrease in exposure of low-income populations (to VTR by both *Aedes* mosquitoes), under all scenario combination except SSP3*RCP8.5 (due to the increase in poverty described under this scenario).

Aggregated at the country and regional level (Figure 5b/c), results clearly show the dominant contribution of the population effect to both increases and decreases in total population exposure to VTR by *Aedes* mosquitoes, with some regional exceptions under SSP3*RCP8.5 (e.g. West and Northeast regions) and *Ae. albopictus*-specific exceptions under SSP5*RCP8.5. This result clearly highlights the crucial role that socioeconomic pathways play in shaping future population exposure to *Aedes*-borne VTR in the United States.

### 3.5. Avoided exposure

The use of the scenario matrix also enables exploring the avoided exposure due to (*i*) shifts in climatic conditions (e.g. resulting from mitigation options) or to (*ii*) shifts in socioeconomic pathways (e.g. resulting from the implementation of different social policies). Aggregated at the national (CONUS) scale (Figure 6), a shift from a high to a low emission scenario (RCP8.5—RCP2.6 shift) leads to a projected decrease in population exposure to *Ae. aegypti* VTR by 20% (*IQR=5.8*) by 2080 (regardless of the population group accounted for), while SSP5—SSP3 and SSP5—SSP1 shifts lead to a higher projected decrease, respectively 52% and 26% (for the total population only). Although results show a dominant effect of demographic/socioeconomic scenarios on avoided exposure, climate mitigation options also play a substantial role in shaping future exposure, particularly in the Northeast and West regions, where a RCP8.5—RCP2.6 shift would lead to greater avoided exposure to *Ae. aegypti* VTR than a SSP5—SSP1 shift.

**Figure 6.**
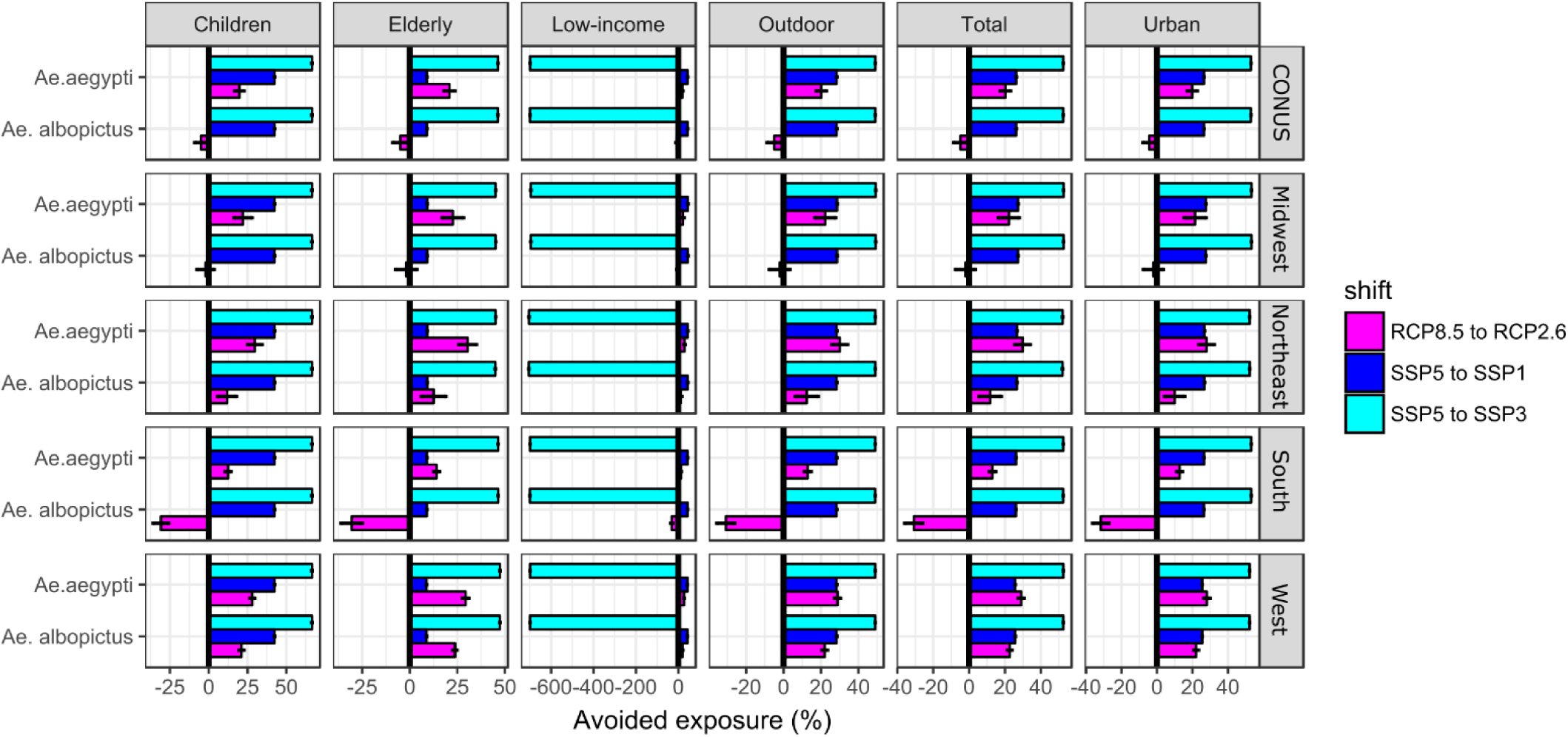
Avoided exposure to *Aedes*-borne VTR, in relative terms (%), due to shifts from RCP8.5 to RCP2.6 (assuming current socioeconomic/demographic conditions and multi-model mean), from SSP5 to SSP1, and from SSP5 to SSP3 (assuming current climatic conditions). Results are shown for year 2080 only and are aggregated at the country (CONUS) and regional level, for the six population groups and the two *Aedes* mosquitoes. Errors bars represent the multi-model interquartile ranges (IQRs).

Regarding exposure to *Ae. albopictus* VTR, shifts in SSPs would lead to avoided exposure of similar magnitude to that of avoided exposure to *Ae. aegypti* VTR, while the effect of a RCP8.5—RCP2.6 shift would be reversed. Indeed, a RCP8.5—RCP2.6 shift would not decrease, but rather increase, exposure to *Ae. albopictus* VTR (by 5% (*7.7*)), highlighting the contrasting influence of climate change scenarios on *Aedes*-borne VTR in the United States. Similar findings apply in the South where a RCP8.5—RCP2.6 shift would increase exposure to *Ae. albopictus* VTR by as much as 31% (*10*). The West and Northeast are the only regions where a RCP8.5—RCP2.6 would decrease population exposure to *Ae. albopictus* VTR (by 29% (*3.3*) and 12% (*11*) respectively).

Trends in avoided exposure for outdoor workers, children, and urban populations follow those of the total population. However, trends differ for the elderly and low-income populations. Avoided exposure of elderly due to a SSP5—SSP3 shift largely dominates the avoided exposure. Conversely, SSP5—SSP1 shift lead to very little avoided exposure, in most cases inferior to the avoided exposure due to RCP8.5—RCP2.6 shifts (for *Ae. aegypti* only). This is explained by the low net difference in the number of elderly between these two scenarios. Finally, due to the large difference in the number of low-income persons between SSP5 and SSP3, a SSP5—SSP3 shift would lead to increased exposure of 700% in all regions and for VTR by both *Aedes* mosquitoes. This highlights again the important contribution of socioeconomic development pathways to future population exposure to *Aedes*-borne VTR in the United-States.

## 4. Conclusions

We projected that population exposure to *Aedes*-borne VTR will increase during the 21^st^ century across the United States, but with contrasting patterns depending on (*i*) the population group of concern, (*ii*) the species of *Aedes*, (*iii*) the emissions scenario (i.e., RCPs), and (*iv*) the socioeconomic pathway (i.e., SSPs). We demonstrated that the type of socioeconomic pathway plays a critical role in shaping future population vulnerability and exposure to *Aedes*-borne VTR, particularly when the pathway projects a decrease in certain vulnerable groups such as low-income populations. Our approach emphasizes the importance of including SSP-based population projections to ensure a more realistic portrayal of future *Aedes*-borne VTR under climate change scenarios. The differential exposure across the myriad SSP-RCP scenario combinations underscores the wide range of potential outcomes (and therefore the need to use scenarios to span future climatic and societal uncertainties), and provides insight into the substantial avoided exposure that certain social policies and mitigation efforts could trigger. One particularly unique aspect of the present study is its breakdown of population projections into potentially vulnerable subgroups. From this, we found that the trends in exposure of some vulnerable subgroups differ from that of the total population. For instance, (*i*) exposure of the urban population increases slightly faster than that of the total population due to the continuing urbanization, (*ii*) exposure of the elderly drastically increases under all SSP-RCP combinations due to the rapid ageing of the US population, and (*iii*) the number of low-income communities exposed to *Aedes*-borne VTR rapidly drops with the decrease of the net low-income population depicted under some scenarios.

While a comprehensive list of limitations is given in Ryan et al. (2019), the most important limitation of the projections of future *Aedes*-borne VTR is the assumption that it is only driven by climate change, when evidence suggests that land use change, urbanization, population growth, migration, and economic development play a significant role in shaping the future transmission of *Aedes*-borne viruses (Messina et al., 2016, Astrom et al., 2012, Alimi et al., 2015, Kraemer et al., 2019). This study is also associated with limitations related to the SSP-based projections of vulnerable population groups (see Text S1), which are highly uncertain. Thus, they are most valuable as means of placing bounds of uncertainty on possible future population outcomes.

We view the SSP*RCP framework as a promising tool to explore the complex interactions among socioeconomic development, climate change, and the future spread of VBDs – as recently highlighted in Messina et al. (2019). The main advantages of this framework include (*i*) the SSPs are being increasingly quantified (on gridded scales) for a number of relevant variables such as population growth (Jones & O’Neill, 2016, Gao, 2017), GDP (Murakami & Yamagata, 2016, Gidden, In review), and urbanization (Gao & O’Neill, 2019, Li et al., 2019), (*ii*) the scenarios account for the wide range of uncertainties in both socioeconomic development type and emission scenarios, (*iii*) the scenario matrix can be used to explore the relative contribution of climate change and socioeconomic development to the future spread of VBDs, and (*iv*) the growing literature on the vulnerability of populations – and of the health sector – under the SSPs (Ebi, 2013, Sellers & Ebi, 2017, Rao et al., 2018, Zimm et al., 2018, Welborn, 2018, Striessnig & Loichinger, 2016) can inform about the future vulnerability of exposed populations.

## Acknowledgments

This work was partly fund by the Swiss National Science Foundation’s Doc Mobility scholarship and by the National Institutes of Health, NIAID R01AI091843. SJR was supported in part by NSF DEB EEID 1518681. The authors declare no known conflict of interest. NCAR is supported by the National Science Foundation. Open access fees were partly covered by the Hub/Institute GEDT at the University of Geneva.

