## Supplementary Material for "Intersecting vulnerabilities: Climatic and demographic contributions to future population exposure to *Aedes*-borne viruses in the United States"

Submitted to *Environmental Research Letters*

Fig S1-S8 p2-7

Table S1-S3 p8-9

Text S1 p10

References p10

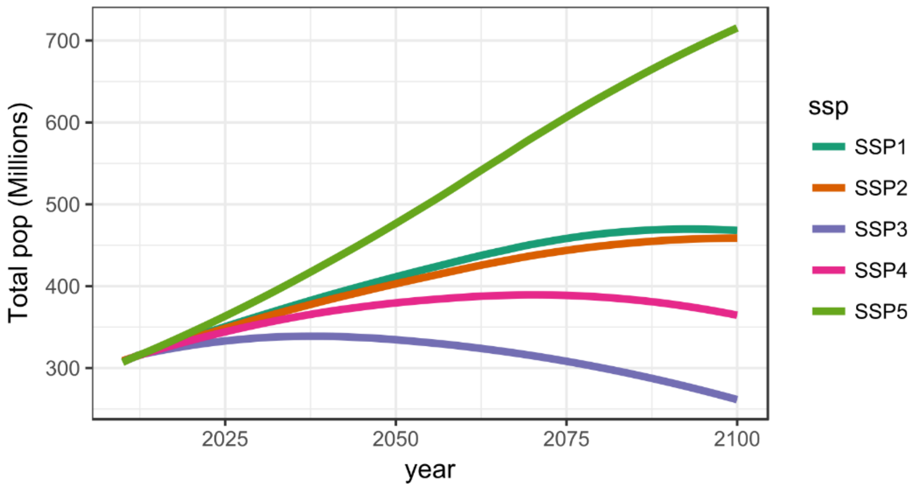


**Figure S1 –** Total population of the United States under the five SSPs, spanning 2010-2100 (KC & Lutz, 2017).

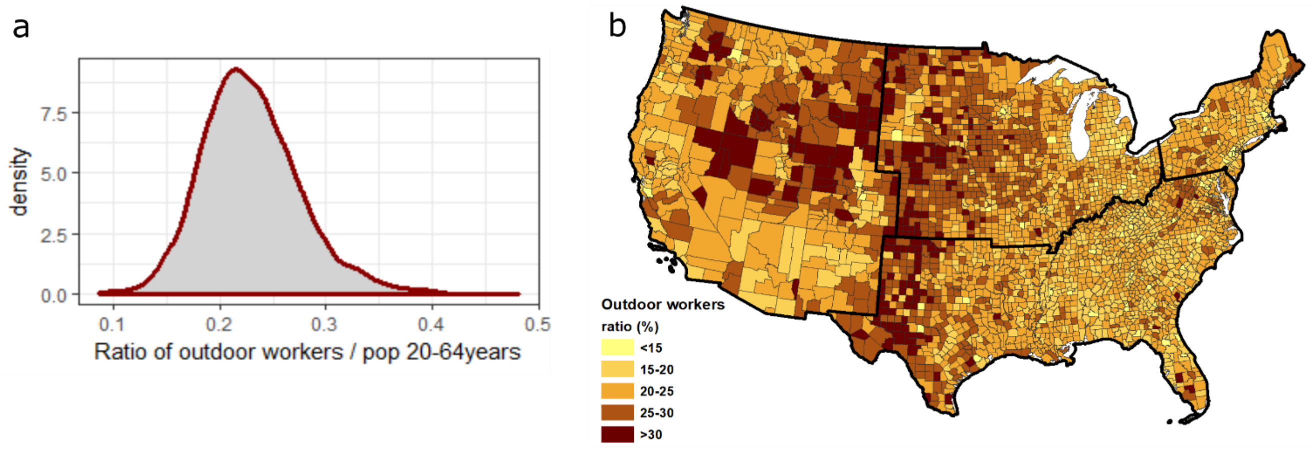


**Figure S2 –** Density plot **(a)** and spatial pattern **(b)** of the county-level ratio of outdoor workers over the working age population (20-64 years), averaged for the period 2010-2017.

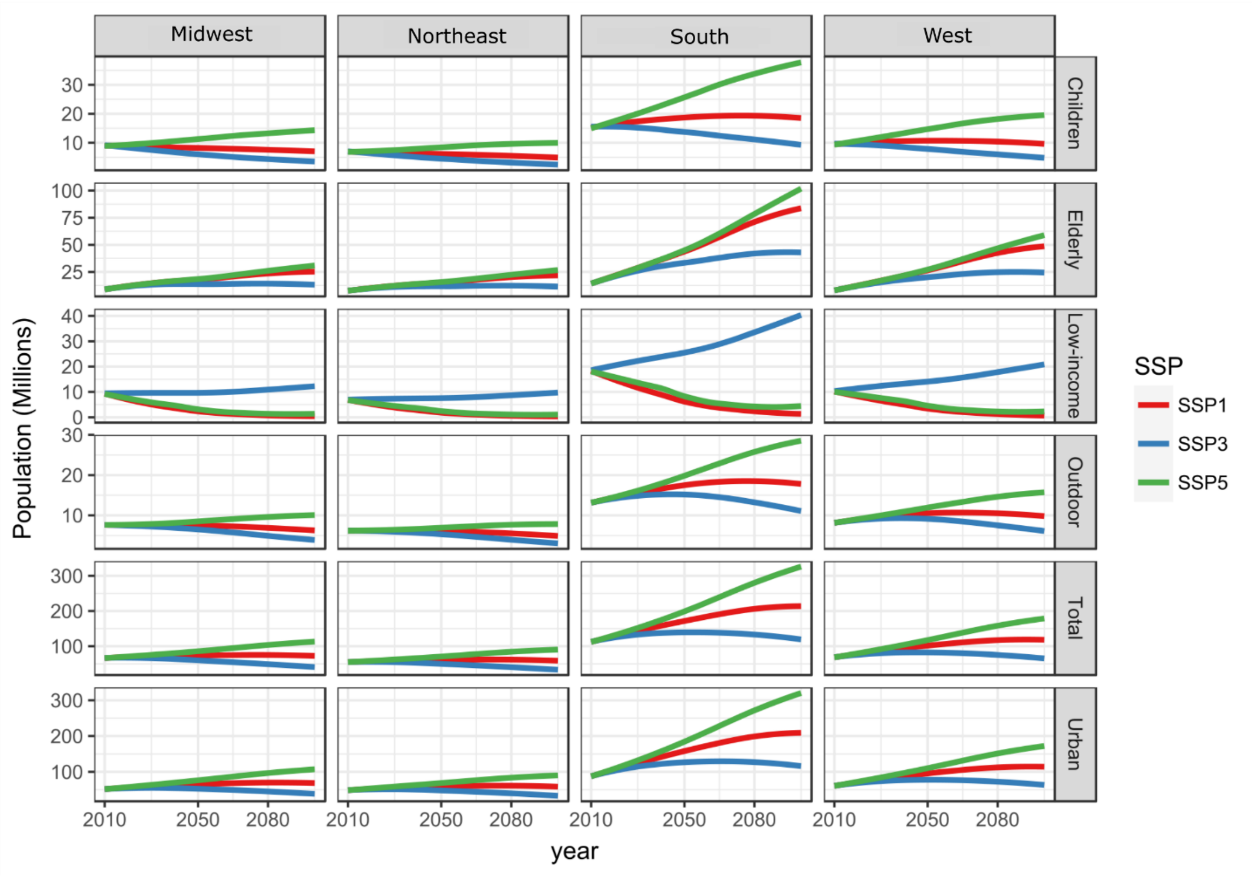


**Figure S3 –** Population projections of the total population and the five vulnerable population groups, under SSP1, SSP3, and SSP5, for the four United-States regions.

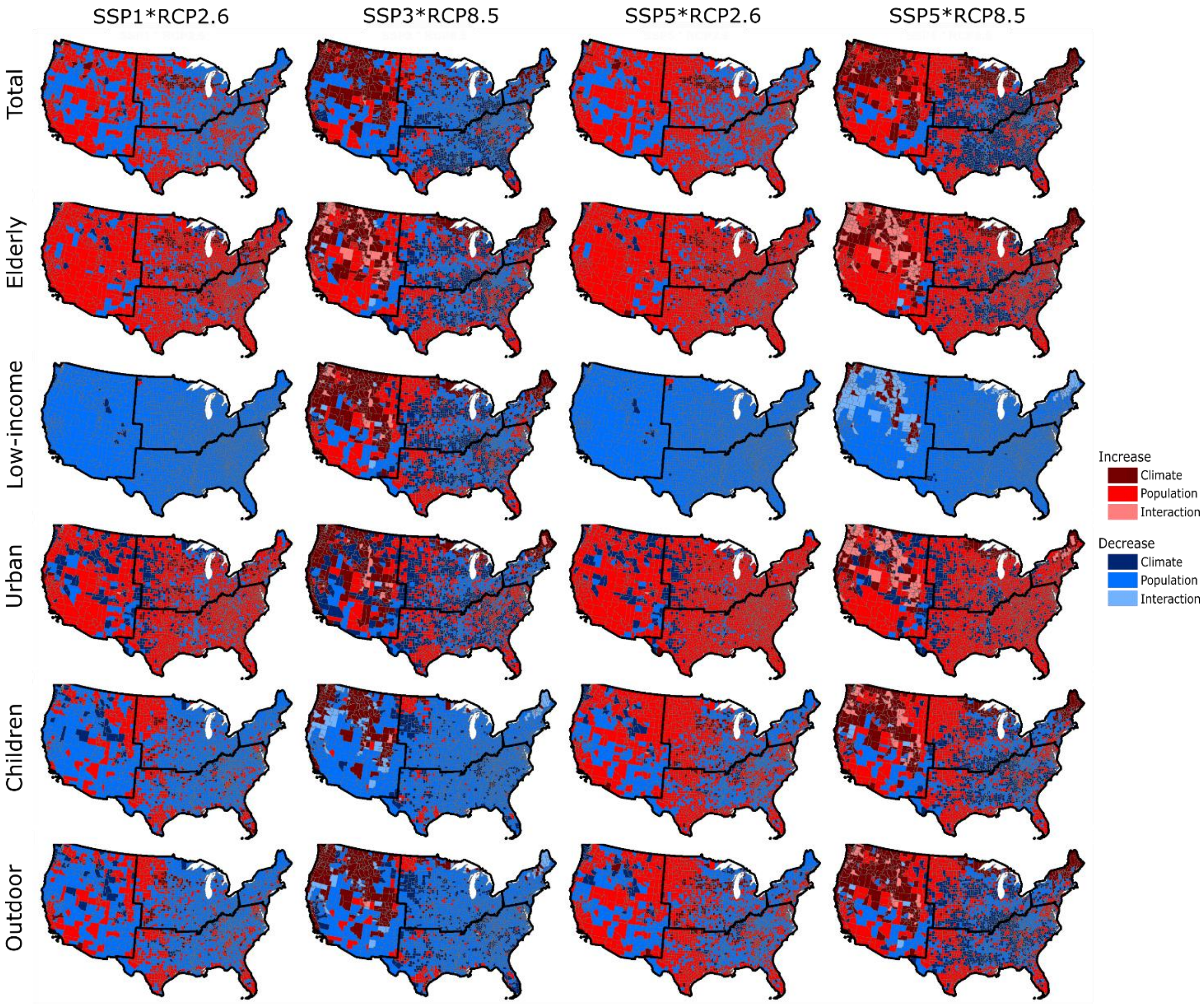


**Figure S4 –** Dominant effect (climate, population, or interaction) responsible for the highest increase (or decrease) in exposure at the county-level, for the six population groups and for exposure to *Ae. albopictus* VTR in year 2080 only, using the multi-model mean.

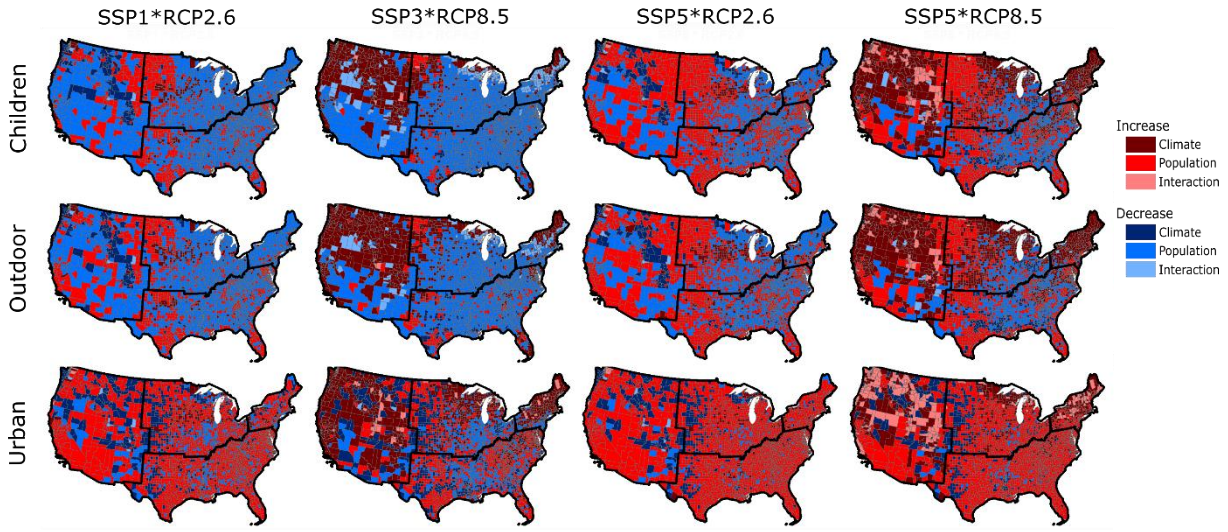


**Figure S5 –** Dominant effect (climate, population, or interaction) responsible for the highest increase (or decrease) in exposure at the county-level, for three population groups and for exposure to *Ae. aegypti* VTR in year 2080 only.

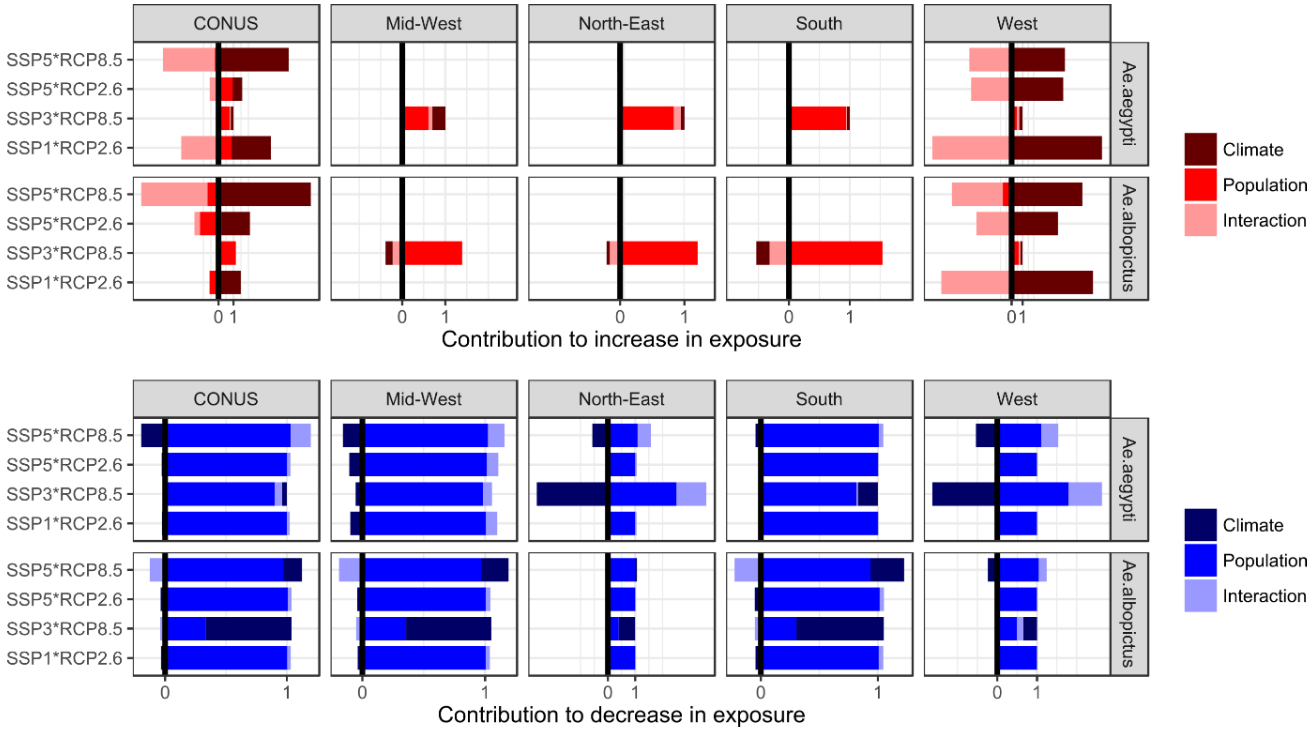


**Figure S6 –** Contribution to increase (top) and decrease (bottom) in low-income population exposure of each individual effect, aggregated at the country (CONUS) and regional scale. Results are presented here for year 2080 only, using the multi-model mean. Note that due to the lack of counties showing an increase in exposure under scenario combinations other than SSP3*RCP8.5 in the South, Northeast, and Midwest region, results for contributions to increase in exposure are showed only for CONUS and West for those scenarios.

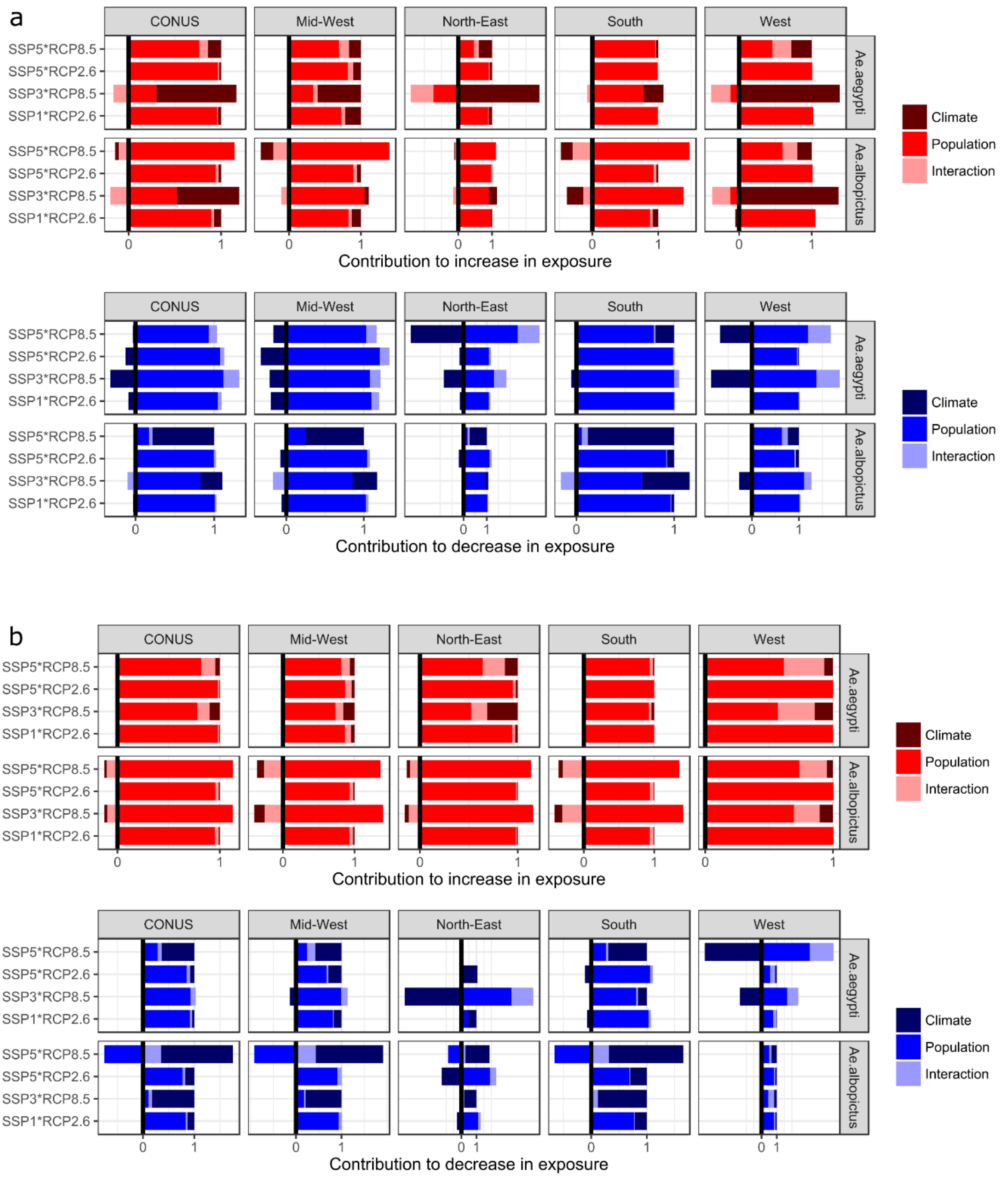


**Figure S7 –** Contribution to increase (top) and decrease (bottom) in children **(a)** and elderly **(b)** population exposure of each individual effect, aggregated at the country (CONUS) and regional scale. Results are presented here for year 2080 only, using the multi-model mean.

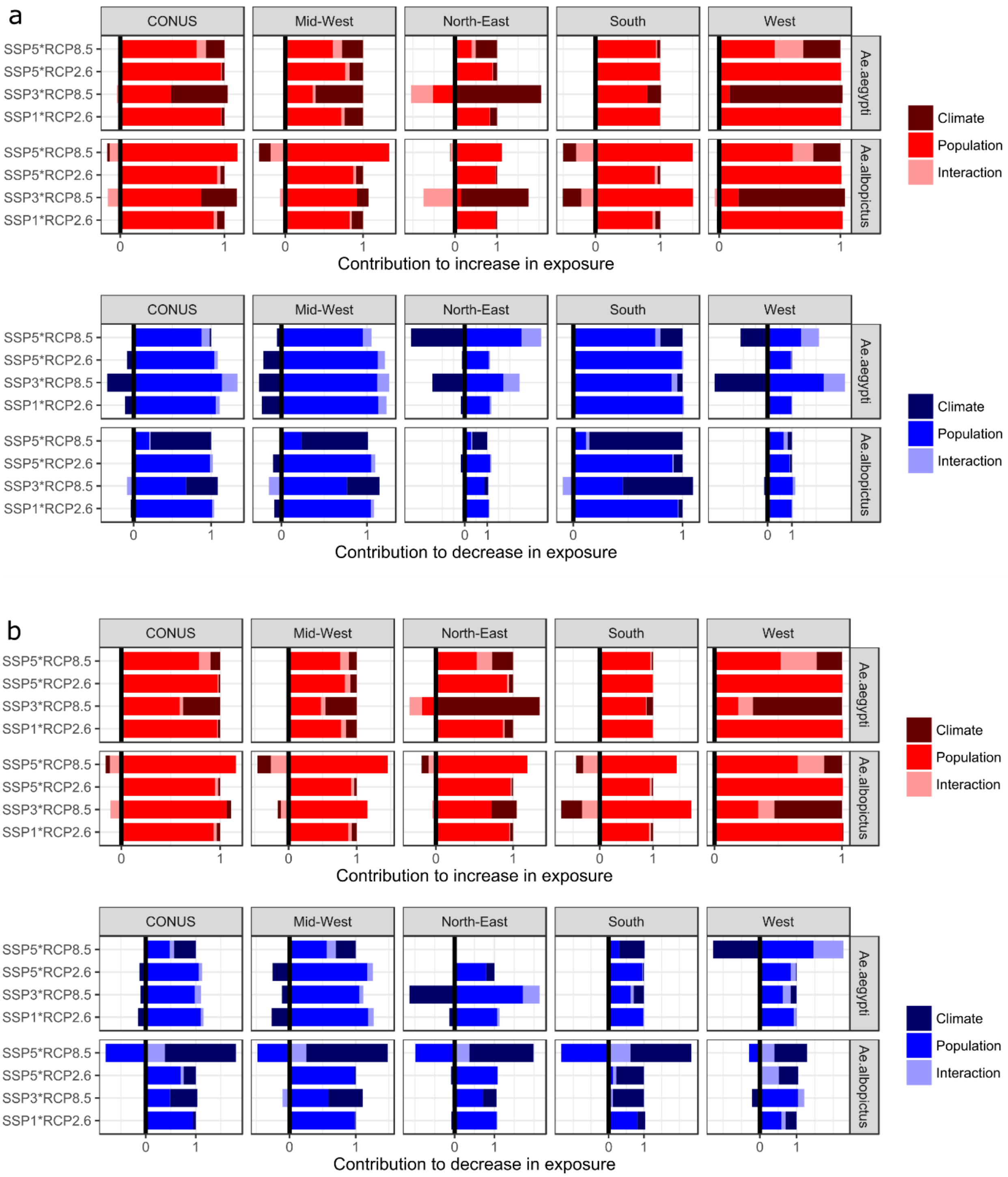


**Figure S8 –** Contribution to increase (top) and decrease (bottom) in outdoor workers **(a)** and urban population **(b)** exposure of each individual effect, aggregated at the country (CONUS) and regional scale. Results are presented here for year 2080 only, using the multi-model mean.

**Table S1 –** List of general circulation models (GCMs) used to project future cumulative monthly transmission risk of *Aedes*-borne viruses (see Ryan et al., 2019).

| Institute & Model | Acronym |
|---|---|
| Beijing Climate Center Climate System Model | BCC-CSM1.1 |
| Hadley Center GCM | HadGEM2-AO |
| Hadley Center GCM | HadGEM2-ES |
| National Center for Atmospheric Research's Community Climate System Model | CCSM4 |

**Table S2 –** Occupations considered as "outdoor workers", with the associated percentage of jobs requiring work outdoors at some point in the day (BLS, 2016). Note that no statistics exist for farming, fishery, and forestry.

| Occupation | Percentage of jobs requiring work outdoors at some point in the day (%) |
|---|---|
| Transportation and material moving | 73 |
| Personal care service | 74 |
| Building and grounds cleaning and maintenance | 77 |
| Installation, maintenance, and repair | 87 |
| Protective service | 91 |
| Construction and extraction | 95 |
| Farming, fishing, and forestry | - |

**Table S3 –** Cumulative monthly transmission risk of *Aedes*-borne viruses, for baseline (2010) and future (2050 and 2080) time-periods, under RCP2.6 and RCP8.5. County-level results are averaged at the regional scale and country (CONUS) scale and separated for *Ae. aegypti* and *Ae. albopictus*.

| *Ae. spp* | Region | Scenario | Year | Mean *(IQR)* | Min *(IQR)* | Max *(IQR)* |
|---|---|---|---|---|---|---|
| *Ae. aegypti* | CONUS | Baseline | 2010 | 2.8 *(0)* | 0.0 *(0)* | 9.4 *(0)* |
| | | RCP2.6 | 2050 | 2.8 *(0.1)* | 0.0 *(0)* | 9.4 *(0.2)* |
| | | | 2080 | 2.9 *(0.3)* | 0.0 *(0)* | 9.3 *(0.4)* |
| | | RCP8.5 | 2050 | 3.5 *(0.3)* | 0.0 *(0)* | 10.1 *(0.7)* |
| | | | 2080 | 4.0 *(0.1)* | 0.0 *(0)* | 11.8 *(0.2)* |
| | Mid-West | Baseline | 2010 | 3.0 *(0)* | 0.0 *(0)* | 5.0 *(0)* |
| | | RCP2.6 | 2050 | 3.0 *(0.2)* | 0.0 *(0)* | 5.0 *(0)* |
| | | | 2080 | 3.1 *(0.3)* | 0.0 *(0)* | 5.0 *(0)* |
| | | RCP8.5 | 2050 | 3.7 *(0.3)* | 0.8 *(0.7)* | 5.0 *(0)* |
| | | | 2080 | 4.1 *(0.1)* | 2.0 *(0.1)* | 5.7 *(0.3)* |
| | North-East | Baseline | 2010 | 1.9 *(0)* | 0.0 *(0)* | 4.0 *(0)* |
| | | RCP2.6 | 2050 | 1.9 *(0.2)* | 0.0 *(0)* | 4.0 *(0)* |
| | | | 2080 | 2.0 *(0.4)* | 0.0 *(0)* | 4.0 *(0)* |
| | | RCP8.5 | 2050 | 2.9 *(0.5)* | 0.0 *(0)* | 4.0 *(0)* |
| | | | 2080 | 3.5 *(0.2)* | 0.0 *(0)* | 5.3 *(0.3)* |
| | South | Baseline | 2010 | 4.8 *(0)* | 1.1 *(0)* | 9.4 *(0)* |
| | | RCP2.6 | 2050 | 4.8 *(0.1)* | 1.0 *(0.3)* | 9.4 *(0.2)* |
| | | | 2080 | 4.9 *(0.3)* | 1.3 *(0.7)* | 9.3 *(0.4)* |
| | | RCP8.5 | 2050 | 5.4 *(0.1)* | 2.3 *(0.2)* | 10.1 *(0.7)* |
| | | | 2080 | 5.7 *(0.3)* | 2.7 *(0.4)* | 11.8 *(0.2)* |
| | West | Baseline | 2010 | 1.4 *(0)* | 0.0 *(0)* | 5.7 *(0)* |
| | | RCP2.6 | 2050 | 1.4 *(0.1)* | 0.0 *(0)* | 5.7 *(0)* |
| | | | 2080 | 1.5 *(0.2)* | 0.0 *(0)* | 5.7 *(0.2)* |
| | | RCP8.5 | 2050 | 2.1 *(0.4)* | 0.0 *(0)* | 5.8 *(0.4)* |
| | | | 2080 | 2.8 *(0.2)* | 0.0 *(0)* | 6.2 *(0.2)* |
| *Ae. albopictus* | CONUS | Baseline | 2010 | 3.1 *(0)* | 0.0 *(0)* | 8.3 *(0)* |
| | | RCP2.6 | 2050 | 3.2 *(0.1)* | 0.0 *(0)* | 11.1 *(0.8)* |
| | | | 2080 | 3.3 *(0.1)* | 0.0 *(0)* | 11.0 *(0.6)* |
| | | RCP8.5 | 2050 | 3.4 *(0.1)* | 0.0 *(0)* | 9.1 *(1.0)* |
| | | | 2080 | 3.4 *(0.2)* | 0.0 *(0)* | 7.8 *(0.2)* |
| | Mid-West | Baseline | 2010 | 3.3 *(0)* | 0.4 *(0)* | 4.6 *(0)* |
| | | RCP2.6 | 2050 | 3.5 *(0.1)* | 0.4 *(0.3)* | 4.9 *(0.1)* |
| | | | 2080 | 3.6 *(0.1)* | 0.4 *(0.4)* | 5.0 *(0)* |
| | | RCP8.5 | 2050 | 3.6 *(0.2)* | 1.9 *(0.2)* | 4.8 *(0.3)* |
| | | | 2080 | 3.5 *(0.3)* | 2.0 *(0)* | 4.8 *(0.3)* |
| | North-East | Baseline | 2010 | 2.7 *(0)* | 0.0 *(0)* | 4.0 *(0)* |
| | | RCP2.6 | 2050 | 2.7 *(0.2)* | 0.0 *(0)* | 4.0 *(0)* |
| | | | 2080 | 2.8 *(0.5)* | 0.0 *(0)* | 4.0 *(0)* |
| | | RCP8.5 | 2050 | 3.4 *(0.4)* | 0.0 *(0)* | 4.6 *(0.2)* |
| | | | 2080 | 3.6 *(0.2)* | 0.0 *(0)* | 4.7 *(0.3)* |
| | South | Baseline | 2010 | 4.4 *(0)* | 2.2 *(0)* | 8.3 *(0)* |
| | | RCP2.6 | 2050 | 4.9 *(0.3)* | 2.2 *(0.2)* | 11.1 *(0.8)* |
| | | | 2080 | 4.8 *(0.4)* | 2.2 *(0.4)* | 11.0 *(0.6)* |
| | | RCP8.5 | 2050 | 4.2 *(0.7)* | 2.3 *(0.3)* | 9.1 *(1.0)* |
| | | | 2080 | 3.8 *(0.4)* | 2.2 *(0.3)* | 7.8 *(0.3)* |
| | West | Baseline | 2010 | 1.8 *(0)* | 0.0 *(0)* | 5.6 *(0)* |
| | | RCP2.6 | 2050 | 1.8 *(0.2)* | 0.0 *(0)* | 5.9 *(0)* |
| | | | 2080 | 1.9 *(0.2)* | 0.0 *(0)* | 5.9 *(0)* |
| | | RCP8.5 | 2050 | 2.5 *(0.3)* | 0.0 *(0)* | 5.9 *(0)* |
| | | | 2080 | 2.9 *(0.1)* | 0.0 *(0)* | 6.2 *(0.4)* |

**Text S1 –** Main caveats of the projections of vulnerable population groups

Outdoor workers projections: By using constant county-level ratio of outdoor workers over the total working age population (20-64 years) of the county population, our projections of outdoor workers account only for demographic changes (population and age-structure). Other changes are likely to influence to future number of outdoor workers at the county level, such as changes in the structure and repartition of the labor force across the different occupations, economic development, technological development, lifestyle changes, etc. Accounting for these changes – which would likely differ across the SSPs – remains challenging and is not the main aim of this study. Furthermore, existing occupational projections are limited to short time-horizon projections, e.g. 2026 for the projections of the Bureau of Labor Statistics (BLS, 2016).

Low-income population projections: Rao et al. (2018) employed three different thresholds of poverty from the World Bank, with the highest threshold being $5.5/cap/day (in $2015 PPP). This threshold is not aligned with the poverty threshold used by the US census bureau (and ACS estimates), which varies between ~$15 and $25/cap/day, depending on the number of persons living in the household. The highest threshold of the World Bank used by Rao et al. (2018) is more representative of extreme poverty in the US than it is of poverty. By aligning our projections of low-income population with the projections of Rao et al. (2018), we assumed the poverty in the US to increase/decrease with the same pace as that of the extreme poverty. This may not be the case. For instance, the large decrease in extreme poverty observed under SSP1 and SSP5 could be associated with only a moderate increase in poverty/low-income communities.

Finally, our projections of the different population groups heavily relies on the county-level population projections depicted in Hauer (2019). Once different sets of county-level population projections for the United-States becomes available, it would be interesting to explore the uncertainty due to population projections, as these have proved significantly influence future population exposure to climate-related hazards in other regions (e.g. Rohat et al., 2019).
